# Differences in nodulation among rhizobia strains can obscure plant benefits per nodule in single-strain experiments

**DOI:** 10.64898/2026.11.16.767161

**Authors:** R. Ford Denison, Nicholas J. Adelmann, Katherine E. Muller, Abraham G. Smith

## Abstract

Each legume plant in the field typically hosts several strains of rhizobia, yet plant benefits from different strains are often compared using single-strain experiments. We present an automated protocol to track nodule growth (using photos and the RootPainter model) and to measure hydrogen production, a nondestructive proxy for nitrogen fixation. Soybean seed yield was highly correlated with hydrogen production (integrated over growth), except in 5 of 20 field isolates of rhizobia, apparent hydrogen recyclers. Most yield differences among strains were due to differences in nodule mass per plant, which might not predict field benefits, rather than benefit per nodule mass. However, three isolates with the greatest nodule mass provided the least yield benefit. Strain rankings were mostly consistent in a second experiment, but two exceptions showed the value of replication. For ten of 12 strains in the second experiment, final seed weight was highly correlated with week-6 estimates of shoot weight, but two exceptions suggest that vegetative growth may not always predict fitness benefits to plants. Although singly-inoculated plants are an imperfect proxy for multiply-infected plants in the field, tracking nodule growth and measuring hydrogen production can make these experiments more informative.

**Highlight:** Nodule data from root photos can reveal whether differences in plant growth with different rhizobia strains are due to more benefit per nodule or only to greater nodulation.

## Introduction

Benefits to plant hosts from different rhizobia strains are usually compared by measuring some combination of plant growth, nodule number or mass per plant, and nitrogen fixation. Because each plant in the field typically hosts multiple strains, mixed-inoculation experiments may be key to predicting field results. However, there is a vast literature of strain comparisons using plants inoculated with single rhizobia strains. We therefore ask what measurements from single-strain experiments would best predict plant benefits and rhizobia fitness when each strain occupies only a fraction of a plant’s nodules.

First, consider relative benefits to plants from different rhizobia strains. In either lab or field, plant benefit from a strain is the product of its nodule number per plant and its net benefit (subtracting costs) per nodule. If differences among strains in plant benefit are partly due to differences in nodule number per plant, is nodule number a confounding factor requiring correction, or an additional plant benefit, complementary to per-nodule benefit? The latter might be true if 1) total nodulation limits legume growth in the field, and 2) nodules/plant in single-strain experiments predicts greater nodulation in the field. However, field experiments have found that a positive yield response to inoculation was rare unless indigenous rhizobia populations were less than 10 cells/g soil, orders of magnitude below values typical of soils where the legume host has been grown recently (Thies et al., 1991). This suggests that strains with greater “nodule-forming efficiency” (McDermott & Graham, 1990) in single-strain experiments would not necessarily increase nodulation in most soils. However, those strains might be relatively competitive for nodule occupancy. If so, that would increase their positive -- or negative -- impact on plants, depending on whether they provide more or less benefit per nodule than those that would otherwise occupy the same nodules.

Next, consider relative fitness benefits to different rhizobia strains from symbiosis with legumes. Fitness benefit to rhizobia depends on the number of rhizobia cells that escape from nodules into the soil and on the resources each rhizobial cell acquires in nodules. These resources include phosphorus (Cassman et al., 1981) and polyhydroxybutyrate (PHB), which can support survival in the soil, perhaps for years (Muller & Denison, 2018). The number of rhizobia cells released into the soil is the product of the number of nodules containing a given strain and the number released per nodule. Although nodule occupancy in the field depends on many factors, strains that form more nodules in single-strain experiments might perhaps be expected to form more nodules in the field. Similarly, although mass per nodule can be an imperfect predictor of rhizobia per nodule (Gubry-Rangin et al., 2010; Sachs et al., 2010; Oono et al., 2011), larger nodules probably release more rhizobia, all else being equal.

However, even actual counts of rhizobia released from nodules of singly-inoculated plants might not reliably predict relative fitness benefits to different rhizobia strains on multiply-infected plants. One reason is the “sanctions” that legumes impose on less-beneficial nodules, reducing nodule growth and rhizobia reproduction (Singleton & Stockinger, 1983; Oono et al., 2011; Westhoek et al., 2021; Porter et al., 2024). Sanctions are based on an individual nodule’s performance (Kiers et al., 2003), not on the plant somehow adding up benefits from all nodules containing that strain. Some legumes even impose within-nodule sanctions (Regus et al., 2017; Daubech et al., 2017). In singly-infected plants, less-beneficial strains may avoid sanctions, because the plant has no better alternative (Westhoek et al., 2021). However, a plant hosting only a less-beneficial strain will have lower rates of nitrogen fixation and photosynthesis (Bethlenfalvay et al., 1978) and therefore fewer resources to support nodules (Gubry-Rangin et al., 2010). With multiply-infected plants, a focal strain’s nitrogen fixation will have less effect on the plant’s overall supply of resources to support nodules, but the plant may impose sanctions on less-beneficial strains that would have escaped sanctions in singly-infected plants (Westhoek et al., 2021). In both singly- and multiply-infected plants, therefore, less-beneficial strains would get less fitness benefit from symbiosis, but differences in mechanism make extrapolation from single-strain experiments risky.

Hormonal manipulation of plants by rhizobia (Denison, 2019; Klein et al., 2022) is another complication. Known mechanisms include rhizobia producing plant-hormone mimics (Tatsukami & Ueda, 2016; Nett et al., 2022), or interfering with a plant’s hormonal signaling (Ruan & Peters, 1992; Belimov et al., 2019). For example, wildtype rhizobia that produce an ethylene inhibitor, rhizobitoxine, gain more PHB per cell. In singly-inoculated plants, this benefit is countered by fewer rhizobia per nodule. When mixed with a rhizobitoxine knockout, however, the wild-type strain gets the PHB benefit without the cells-per-nodule cost (Ratcliff & Denison, 2009). This example shows that experiments to predict the fitness of rhizobia strains in the field should use multiply-infected plants.

Rather than relying on a single variable, medical professionals use multiple “vital signs” (temperature, blood pressure, pulse) to assess patient health. We therefore developed efficient, nondestructive methods to track nodule number, nodule mass, and plant growth over weeks, using photography and RootPainter software (Smith et al., 2022). We also track hydrogen production, a byproduct of nitrogen fixation (Layzell et al., 1984). The hydrogen assay is not useful for the minority of rhizobia strains (Keyser et al., 1984) that recycle hydrogen. With our automated approach, getting all these data on 70 plants at least weekly throughout growth requires about 16 hours of undergraduate-student work per week, much of it for thrice-weekly watering by weight, which also generated evapotranspiration data.

We tested the hypothesis that rhizobia strains differ mainly in specific nodule activity (nitrogen-fixation rate per nodule mass), perhaps corrected for carbon cost. This “SNA hypothesis” assumes that strains differ little in nodulation. If this is true, then strain rankings based on whole-plant measurements could also correctly rank strains by benefits per nodule. Benefits per nodule are key to plant sanctions and to the increase in crop yield for a given increase in nodule occupancy by a focal strain. If, instead, there are major differences in nodulation among strains, then one would need to correct for nodulation differences to estimate per-nodule benefits. Furthermore, if strains have large differences in the timing of nitrogen fixation, then strain rankings based on early vegetative growth might poorly predict effects on plant fitness or seed yield.

Previous research has shown that the “SNA hypothesis” is not always true. For example some plant-benefit ranks for rhizobia strains based on clover biomass reversed between 3 and 6 months (Briggs et al., 2026). Many studies lack the data needed to test the hypothesis for particular species or conditions, however, because counting nodules or measuring nodule mass is labor-intensive. Furthermore, destructive assays over plant growth require large numbers of plants, limiting the numbers of strains or treatments that can be tested.

We therefore automated nondestructive acquisition of nodule data with computer-controlled photography (during the watering and weighing operation) and analyzed the resulting photos with RootPainter. This protocol makes taking these data routine, at least for experiments in growth pouches. We used 20 field isolates of rhizobia to illustrate our approach and to test the “SNA hypothesis.”

## Materials and Methods

Briefly, we inoculated soybean plants with one of 20 different field isolates, grew plants to maturity, then weighed shoots and seeds. Throughout growth, we measured evapotranspiration and hydrogen production. Photos of nodulated roots and shoots were used to track growth. We repeated the experiment with a subset of the 20 isolates, to check the consistency of strain rankings for nodulation and plant benefit.

Rhizobia were isolated using trap plants in soil from plots with a 30-year history of either maize or soybeans every year. From each host-availability treatment, we chose five isolates with high and five with low PHB production in nodules, measured by flow cytometry (Muller & Denison, 2024).

Rhizobia isolates (putative “strains”) were maintained at −80 C in arabinose-gluconate (AG) medium (Sadowsky et al., 1987) + 25% glycerol, then streaked on AG plates and grown in liquid AG medium. Growth pouches 16.5 cm wide x 35 cm long (Mega International, Roseville, MN) were inoculated with approximately 3 million cells each (estimated from optical density and previous plate counts) in 50 mL of Fujikake’s N-free plant-growth medium (Fujikake et al., 2002), supplemented with 20 mg NH_4_NO_3_/L (10 mg/L in the repeat experiment), to mimic a soil supplying some nitrogen but not enough to suppress nodulation. One seed of dwarf soybean cultivar “MN0095” (surface sterilized in 3% hydrogen peroxide) was added to each pouch, along with a plastic straw. We started 10 replicate plants for each of the 20 rhizobia strains, then blindly culled pouches with delayed emergence or very poor growth, prior to nodulation by any strain. This resulted in 5 to 7 replicate plants per rhizobia strain.

Plants were grown in a Conviron E7/2 growth chamber with 14 h light at 22 C and 18 C at night. Growth pouches were watered through the straws (to limit cross contamination) three times per week with the Fujikake+NH_4_NO_3_ solution. Water use was measured using an Ohaus Scout balance connected to a Raspberry Pi 4, which also photographed the shoot and the nodulated root, backlit with a white LED panel. Growth pouches were drained and refilled with nutrient solution once per week. At seed maturity, plants were air dried and weights of shoots (minus lost leaves) and seeds were recorded.

We measured hydrogen production by nitrogenase as a proxy for nitrogen fixation by attached nodulated roots. The proportional relation between hydrogen production and nitrogen fixation (both in air) is set by the Electron Allocation Coefficient (EAC) between production of ammonia versus hydrogen (Layzell et al., 1984). We previously found that the greater N-per-C efficiency of PHB-minus knockouts were not due to differences in EAC (Oono et al., 2020). We did not measure EAC or overall efficiency in these experiments. Rubber gaskets were used to seal growth pouches and connect them to our gas-analysis system. Pouches received room air pumped at 500 mL/min through a bubbling humidifier, while gas from the pouch returned through a water trap at 300 mL/min to a Figaro TGS2600 hydrogen sensor, with the excess vented from pouch (Figure 1A). Solenoid valves (not shown in Figure 1A) were added to speed switching between plants.

**Figure 1.**
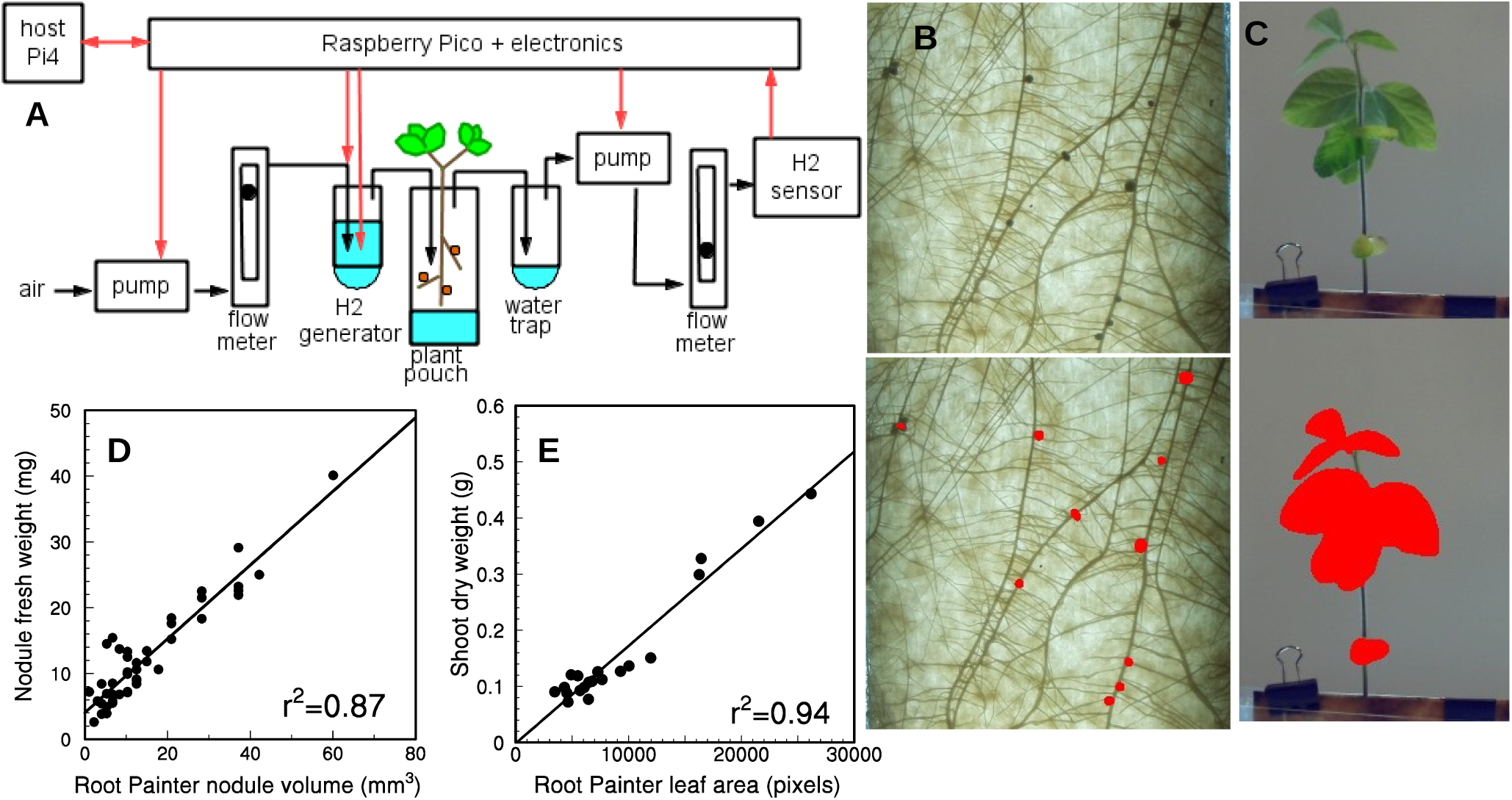
Hydrogen production was measured as a proxy for nitrogen fixation, whereas fresh weights of nodules and dry weights of shoots were estimated by regressing harvested weights on estimates from Root Painter analysis of photos. A) Air flowed through a humidifying hydrogen generator and plant-growth pouch to the hydrogen sensor, controlled by a Raspberry Pi Pico and a Raspberry Pi host computer. B) Original root image above image with red-labeled nodules identified by Root Painter model. C) Original shoot image above image labeled by Root Painter trained on leaves. D) Measured fresh weights of harvested nodules versus Root-Painter estimates of each nodule’s volume. E) Relation between Root Painter estimates of projected leaf area and measured dry weights of harvested shoots. ALT TEXT: Figure 1. Hydrogen production was measured as a proxy for nitrogen fixation, whereas nodule mass and projected leaf area were estimated from photos using the Root Painter software. A) Air flows through a pump, a flowmeter, a humidifying hydrogen generator, a growth pouch with nodulated plant, a water trap, and another pump and flowmeter to the hydrogen sensor, all controlled by a Raspberry Pi Pico and a Raspberry Pi host computer. B) A root with about 15 nodules and one false positive labeled by Root Painter. C) A soybean root photo and the same image with leaves and cotyledon labeled by Root Painter. D) Graph showing linear relation between Root Painter estimates of nodule volumes and measured fresh weights, with r-squared = 0.87. E) Graph showing linear relation between Root Painter estimates of projected leaf area and dry weights of harvested shoots.

The sensitivity of the sensor to hydrogen depends on humidity, so we calibrated the hydrogen sensor at high humidity (from water in the hydrogen generator and the wet paper wick in a growth pouch without a plant) using hydrogen generated by electrolysis. This was an alternative to using magnesium perchlorate to dry the sampled gas (Layzell et al., 1984). Hydrogen production in air was measured repeatedly for each plant during growth. In the first experiment, we used the same hydrogen-calibration curves for all assays, but our software zeroed the hydrogen-sensor reading (to 150 mV, to avoid negative voltages to the Raspberry Pi Pico) before each day’s assays. In the second experiment, with an improved board design, we did a four-point calibration of each board before each set of assays.

Differences among strains in nitrogen fixation per plant could be due to differences in activity per gram nodule or to differences in nodule mass per plant. To distinguish between these possibilities, we estimated nodule mass per plant by applying RootPainter software (Smith et al., 2022), to photos of nodulated roots. After training, RootPainter identified most nodules, with a few false positives and negatives (Figure 1B). Measured fresh weights of nodules were highly correlated with volumes calculated (assuming spherical nodules) using diameter estimates from RootPainter (Figure 1D).

However, some small nodules were heavier than predicted, while large nodules tended to be lighter than predicted. Nodule volumes were therefore corrected using the regression equation: fresh weight (mg) = 4.04 + 0.56 * RootPainter estimate (mm^3^). Between the first and second experiments, we trained RootPainter with shoot photos, with equally promising results (Figure 1C, E).

### Statistical analysis

In some cases we fit a line to a subset of isolates (e.g., excluding putative hydrogen recyclers) to explore an apparent relationship. However, some isolates could be so closely related that they cannot be considered independent replicates. Therefore, apparent regression coefficients should not be considered definitive. Statistical analyses that assume independence (such as correlations between plant benefit and PHB or hydrogen recycling) will be postponed to a second paper, pending phylogenetic analysis.

## Results

Hydrogen production in air is shown for four representative strains in Figure 2A. Values below zero in week 11 could be due to drift in sensor response in weeks since calibration. Labor shortages resulted in some plants being skipped some weeks, but each data point represents at least two replicate plants. Of the four strains, strain I produced the most hydrogen at week 4, but it was soon surpassed by strains A and G. Based on these data alone, we would conclude either that strain M fixes very little nitrogen, or that it is a hydrogen recycler, releasing hydrogen only when its hydrogen production exceeded its recycling capability.

**Figure 2.**
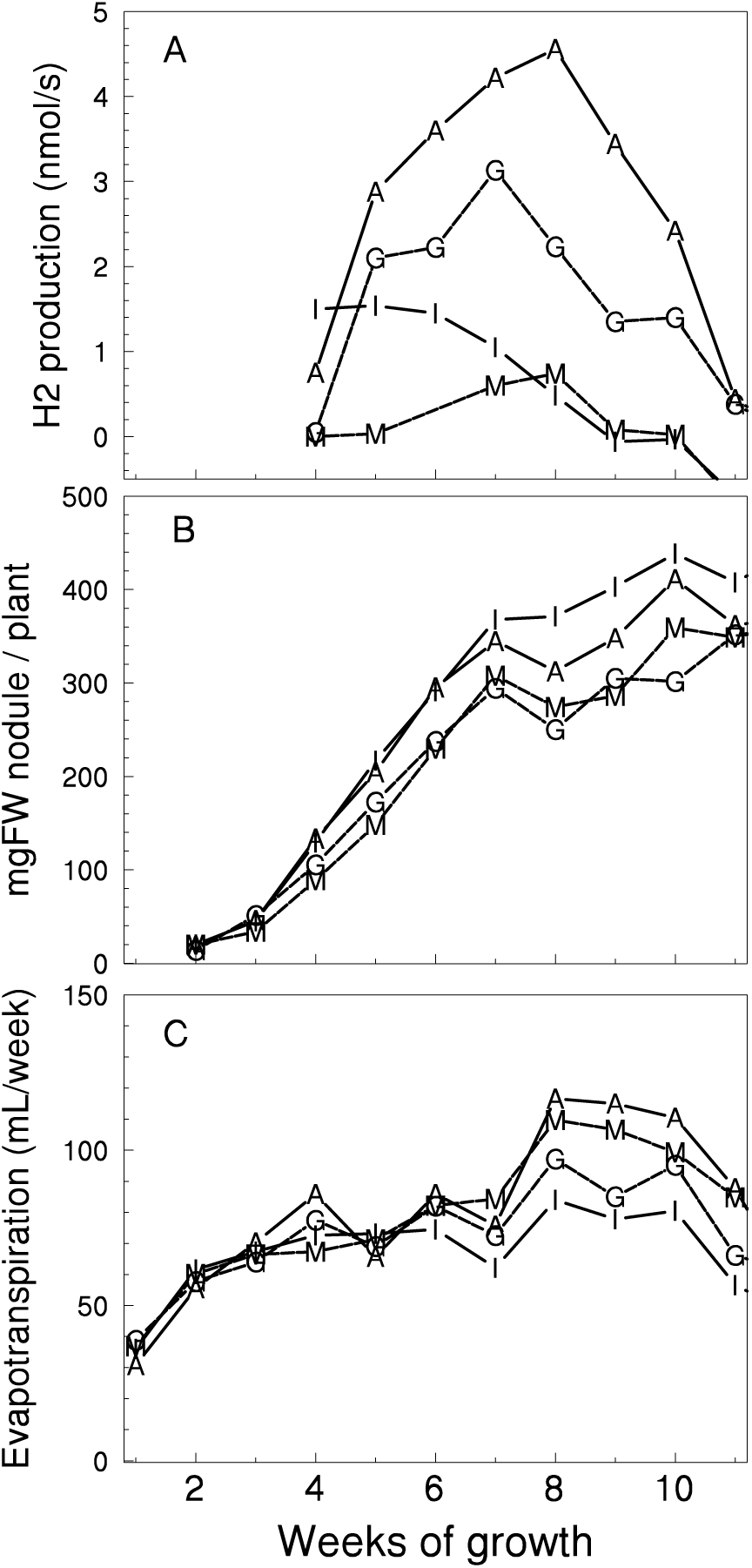
Trends in hydrogen production, nodule fresh weight per plant, and evapotranspiration over weeks for plants inoculated with four example strains of rhizobia, identified by letters. A) Hydrogen production (measured concentration times flow rate). Before week 4, only a few preliminary measurements were made. B) Root Painter estimates of nodule fresh weight per plant, corrected using Figure 1D regression. C) Evapotranspiration, based on weight of water added to per week to maintain depth in pouch. ALT TEXT: Figure 2. Trends in hydrogen production, nodule fresh weight per plant, and evapotranspiration over weeks for plants inoculated with four example strains of rhizobia. A) Hydrogen release by strains A and G peak around week 8. Strain I decreases from week 5. Strain M is near zero most weeks. B)Nodule mass per plant increases over weeks and is greatest for strain I, but differences among strains are less than for hydrogen. C) Evapotranspiration increases to week 8 then decreases and is lowest for strain I.

Estimated nodule fresh weights per plant are shown for the same four strains in Figure 2B. Nodule mass per plant continued to increase somewhat after week 8, even as hydrogen production decreased. Strain I produced the most nodule mass, despite releasing little hydrogen (Figure 2A, B). Figure 2C shows evapotranspiration by growth pouches with plants inoculated with the same four strains. Evaporation from the wet paper wick could be a significant fraction of the total, especially early in growth. To the extent that greater evapotranspiration is due to greater area of transpiring leaves, strain I had the least leaf growth, consistent with its low nodule activity in Figure 2A, whereas strain M’s apparently high leaf growth is consistent with Figure 2A only if its low hydrogen release is due to hydrogen recycling.

Seed yields for soybean plants inoculated with each of our 20 rhizobia strains are shown in Figure 3. In figure 3A, the independent variable is total hydrogen production, estimated by trapezoidal integration over weeks of curves like those in Fig. 2A. Strain M was one of four that produced high seed yields, despite releasing very little hydrogen. We conclude that these strains recycle hydrogen, making their hydrogen release a noninformative proxy for their nitrogen fixation. For 15 other strains, there was a strong correlation (r^2^=0.75) between total hydrogen production and seed yield. Strain Q supported more seed yield than expected from its hydrogen production, but less than the other hydrogen recyclers.

**Figure 3.**
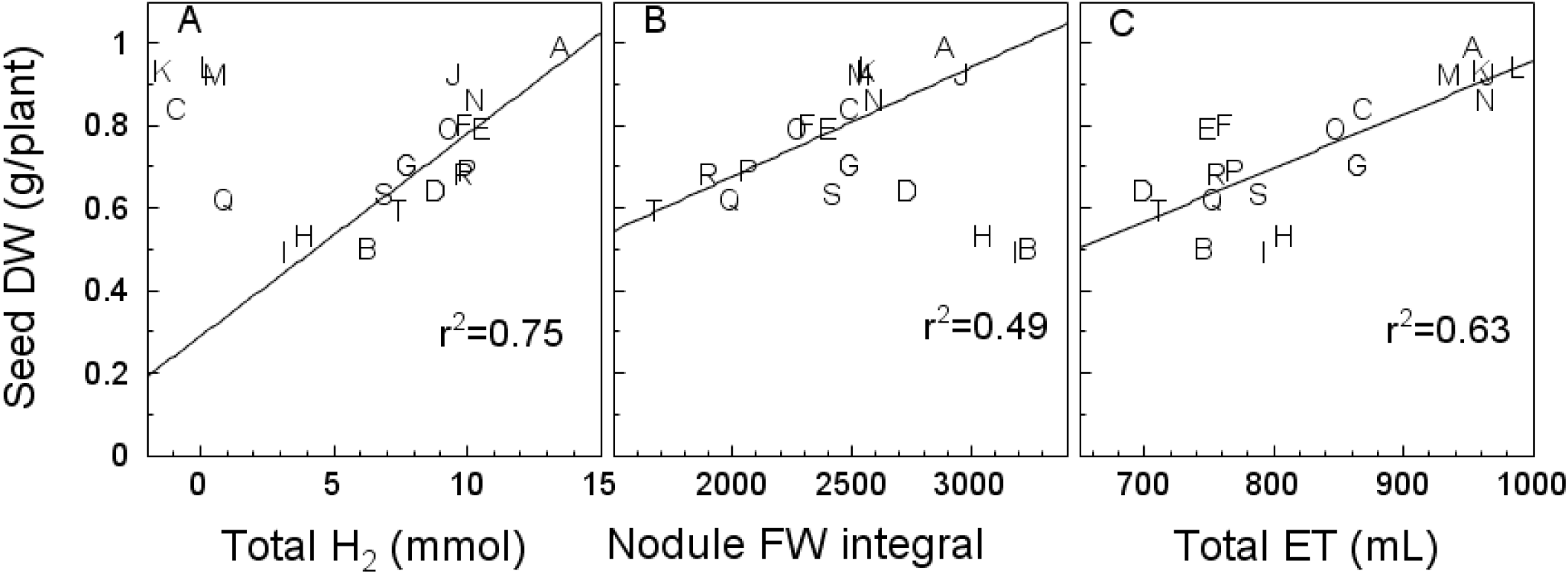
Mean weight of harvested seed for plants, each inoculated with one of 20 rhizobia isolates (identified by letters) was correlated with trapezoidal integrals over weeks of hydrogen production, estimated nodule fresh weight per plant, and total evapotranspiration. A) Seed dry weight versus total hydrogen production over growth by integration of curves like those in Figure 2A, assuming zero fixation in week 3 and earlier. Regression line excludes strains, assumed to be hydrogen recyclers, whose plants released less than 2 mmol total hydrogen. B) Seed weight versus integrals of nodule mass curves like Figure 2B. Regression excludes strains H, I, and B, to emphasize that most strains have similar seed-yield benefits, relative to their nodule-mass integral over weeks. C) Total evapotranspiration over growth and regression line for all strains. ALT TEXT: Figure 3. Mean weight (by rhizobia strain) of harvested seed for plants, each inoculated with one of 20 rhizobia isolates was correlated with trapezoidal integrals over weeks of hydrogen production, estimated nodule fresh weight per plant, and total evapotranspiration. A) Five strains released little hydrogen, although four of them supported among the highest seed yields. Excluding those putative hydrogen recyclers, there was a strong correlation (r-squared = 0.75) between seed yield per plant and the integral of hydrogen production over growth. B) Excluding three low-benefit strains (B, H, and I), seed yield (which ranged from 0.6 gram/plant for strain T to 1 g/plant for strain A) was correlated (r-squared = 0.49) with the integral of nodule fresh weight over growth. C) Seed yield was correlated (r-squared = 0.63) with total evapotranspiration.

Figure 3B plots seed yield against the integral over weeks of estimated nodule mass per plant, based on RootPainter analysis of root images. The “SNA hypothesis” predicts that differences in plant benefit among strains only reflect differences in benefit per nodule mass. To test this hypothesis, we excluded outlier strains B, H, and I, then regressed seed yield on nodule mass per plant. Rhizobia strains A and T differed almost twofold in seed yield, but this was almost entirely due to a difference in nodule mass per plant, not in yield benefit per nodule mass. In contrast, strains B, H, and I produced less seed-yield benefit than any other strain, despite having greater nodule-mass integrals.

Plants inoculated with strains B, H, and I apparently transpired less water, relative to the average for all 20 strains (Figure 3C), but strains E and F supported about 60% more seed yield than those, despite similar water use. Much of the difference in water use among strains is presumably due to differences in plant size, especially the area of transpiring leaves, but we cannot exclude hormonal effects on stomata. The leaf-area assay added between the first and second experiments will facilitate testing such hypotheses.

We also compared strain performance in the two experiments by three measures: seed yield per plant, seed yield per final nodule mass, and seed yield per final nodule number (Figure 4). Relative effects of the different rhizobia strains on seed yields were similar between years, but with some changes in rank order (Figure 4A). In particular, yield with strain T was about half that with strain A in 2025, but approached that of strain A in 2026. In both years, however, these two strains had almost identical yield benefit per nodule mass (Figure 4B), consistent with Figure 3B, or per nodule (Figure 4C). On a benefit-per-nodule basis (Figure 4B, C), strains I, H, and B formed a separate group, inferior to all the other strains.

**Figure 4.**
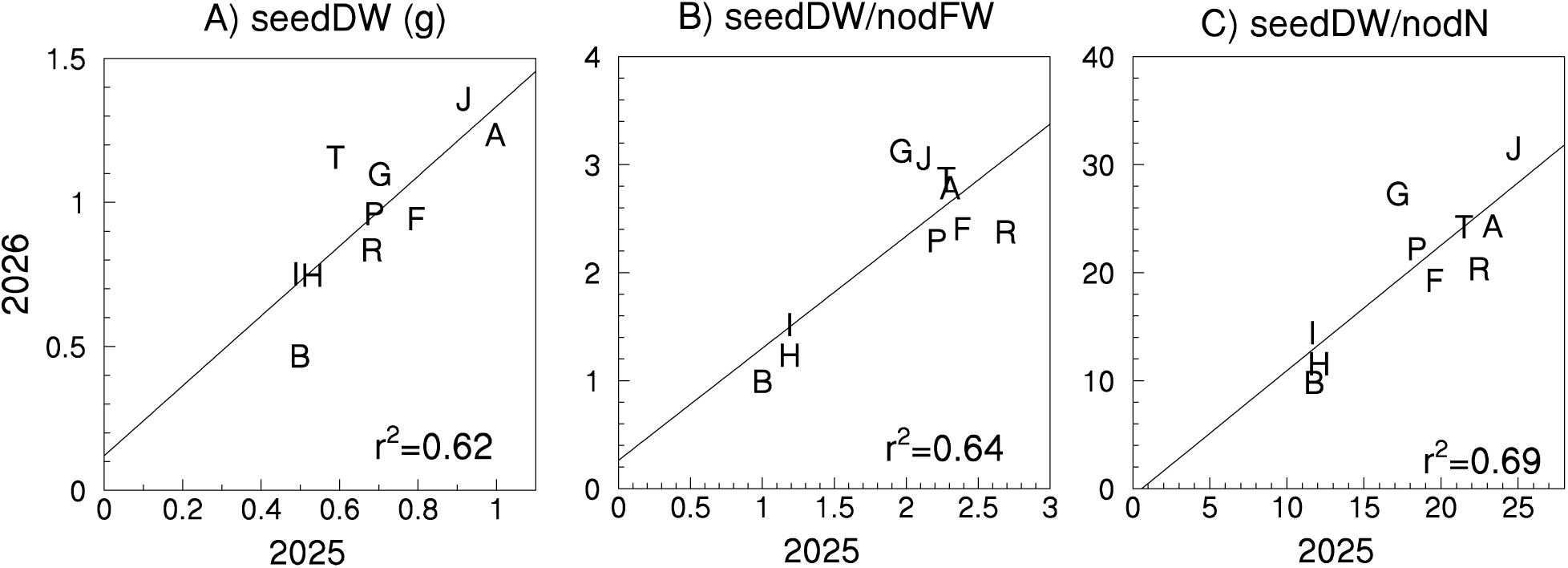
Comparison of 2025 versus 2025 experiments, excluding putative hydrogen-recycling strains (not used in 2026). A) Total seed dry weight per plant in 2026 versus 2025. Strains A and T were more similar in 2026 than 2025. B) Ratio of seed dry weight to final nodule fresh weight in 2026 versus 2025. Note overlap of strains A and T and clustering into two groups, relative to Figure 4A. C) Ratio of seed dry weight to final number of nodules per plant. ALT TEXT: Figure 4. Comparison of 2025 versus 2025 experiments, excluding putative hydrogen-recycling strains (not used in 2026). Correlation (r-squared) between 2025 and 2026 values for seed yield, the ratio of seed yield to nodule fresh weight, and the ratio of seed yield to nodule number per plant was 0.62, 0.64, and 0.69, respectively. Low-benefit strains B, H, and I form a separate cluster from the other strains, especially in the two graphs corrected for nodule fresh weight or number.

The “SNA hypothesis” predicts that strain rankings for plant benefit during vegetative growth should be highly correlated with rankings for seed yield. This was mostly true in our followup experiment, which excluded the putative hydrogen-recycling strains, but added nonfixing (Hahn & Studer, 1986) and a dsRed-labeled (Hayashi et al., 2014) mutants of USDA110, a hydrogen recycler (Figure 5A).

**Figure 5.**
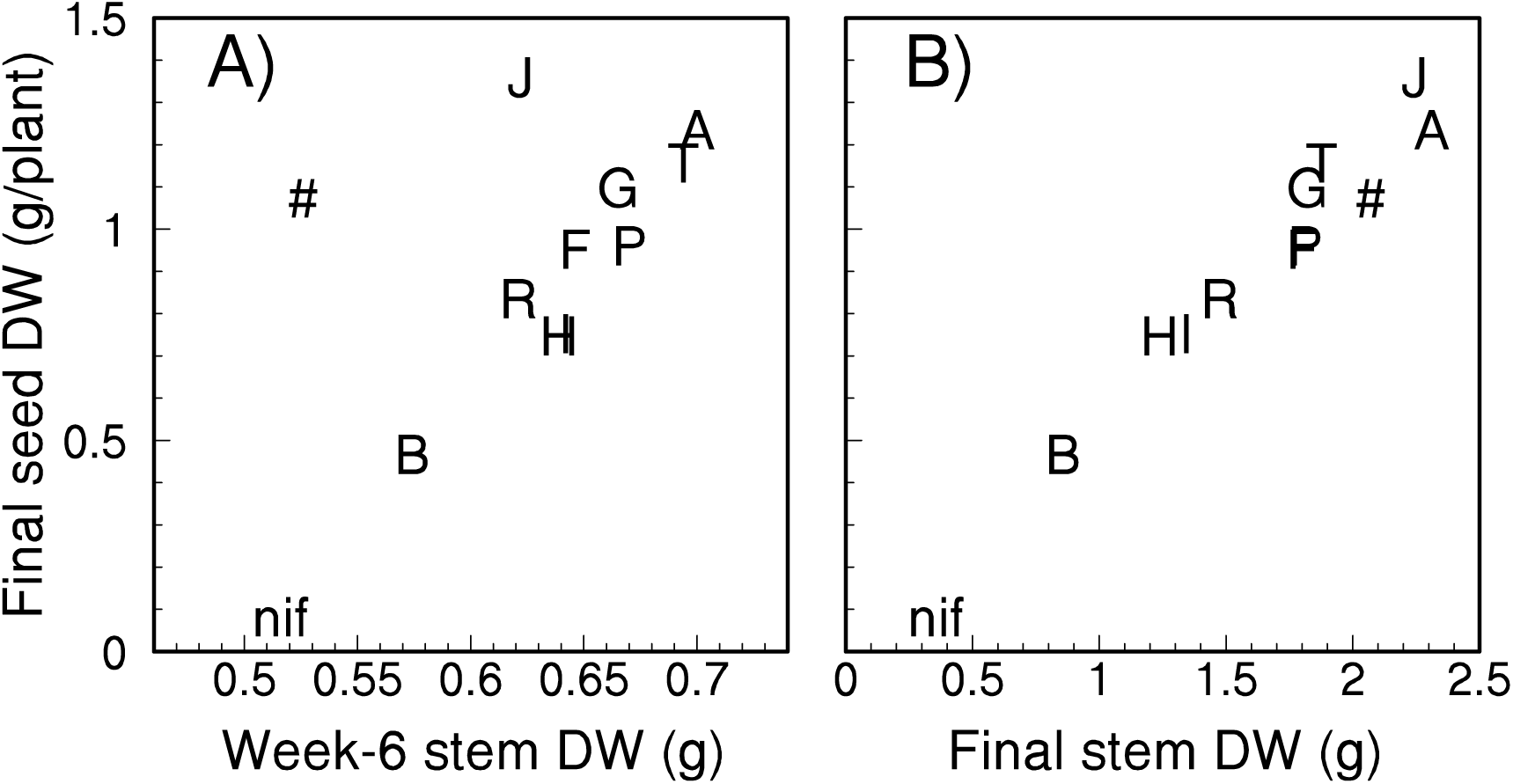
Final seed dry weight was mostly correlated with the estimated or measured shoot dry weights for the same rhizobia strain. # is a dsRed mutant of USDA110. A) Shoot dry weights for week 6 were estimated from shoot photos (see Fig. 1E). B) Shoot and seed dry weights for plants harvested after week 12 were measured after air drying and averaged by rhizobia strain. AL TEXT: Figure 5. Final seed dry weight was mostly correlated with the estimated or measured shoot dry weights for the same rhizobia strain. A) Strain J and the dsRed mutant both had greater seed yield than expected from their estimated week-6 shoot biomass. All other strains, including a nonfixing mutant with near-zero seed yield, fell on a line. B) The relationship between seed yield and final shoot weight was linear, with no significant outliers.

Strain J and the dsRed strain both supported greater seed yield than predicted from week-6 estimates of stem dry weight based on RootPainter’s estimates of leaf area and its correlation with shoot mass (Figure 1E). The discrepancies for these two strains could be due to errors in leaf-area estimates (e.g., due to leaf overlap) or in the relationship between leaf area and stem dry weight, which was not considered reliable once pods formed. Final shoot weights, however, were strongly correlated with seed weights over a five-fold range (Figure 5B).

## Discussion

Five of 20 field isolates supported much more seed yield than expected based on their measured hydrogen production, relative to the other 15 strains (Fig. 3A). These seed yields presumably required high rates of nitrogen fixation, so we conclude that these strains are probably Hup^+^ hydrogen recyclers. The high seed yields of the putative Hup^+^ strains is consistent with this rhizobia trait being beneficial to plants. However, the relatively low (25%) frequency of the trait in our sample suggests that having hydrogen-recycling capability is not particularly beneficial to rhizobia fitness, either directly (by supporting reproduction or PHB synthesis) or indirectly (by reducing the risk of host sanctions). A twelve-state survey found that only 20% of soybean-nodulating rhizobia were Hup^+^ (Keyser et al., 1984). Perhaps a fitness benefit of hydrogen recycling to rhizobia in nodules is balanced by some fitness cost during their sojourn in soil.

The high correlation between seed yield and measured hydrogen production by our 15 putative Hup^-^ rhizobia isolates (Figure 3A) is consistent with the hypotheses that: 1) differences in seed yield are mainly due to differences in nitrogen fixation among strains, 2) the relationship between nitrogen fixation and hydrogen production (Electron Allocation Coefficient) varies little among our rhizobia strains, consistent with previous results for bean (Oono et al., 2020), and 3) hydrogen measurements by our instrument have a consistent proportional relationship to hydrogen production by nodulated roots in growth pouches. The last of these hypotheses, if true, would justify our practice of calibrating the hydrogen sensor in humid air returning from growth pouches, rather than drying the sampled gas, as others have done (Layzell et al., 1984).

These three hypotheses are mutually consistent, in the sense that changing one of them would require a balancing change in another to maintain the high correlation in Figure 3A. For example, what if strain A provided some large hormonal benefit to plants, in addition to fixing nitrogen? Then its ratio of seed yield to hydrogen production would only match that of other strains along the regression line in Figure 3A if its hydrogen production overestimated nitrogen fixation (i.e., lower EAC than other strains). Such balancing of departures from the above hypotheses seems unlikely to be consistent enough to explain Figure 3A. We don’t exclude the possibility of small benefits or costs unrelated to nitrogen fixation, even among our 20 strains. However, Figure 3A is consistent with the very strong genetic correlation between the seed yield and fixed nitrogen of different pea cultivars and their total nitrogen fixation (Millar et al., 2026).

What EAC values are implied by these data? Figure 3A seems to show that a plant releasing 15 mmol H2 would make about 1 g of seeds. Soybean grain is about 6.8% nitrogen (Connor et al., 2011), so the seeds would contain 68 mg N or 4.8 mmol N, equivalent to 2.4 mmol N2 or 14.4 mmol electrons.

Making 15 mmol H2 would require 30 mmol electrons. EAC is the fraction of total electrons going to N fixation, so these data imply an EAC of only 32% (14.4/44.4). This is below the reported range of 40-60% (Schubert & Evans, 1976). However, additional N in stem, roots, or lost leaves would imply more N fixation and thus a higher EAC. Furthermore, the estimate of 15 mmol H2 is based on the assumption that measured hydrogen-production rates continue throughout the day, rather than decreasing at night.

On the other hand, some of the seed N presumably comes from the NH_4_NO_3_ included in our nutrient solution. Isotope experiments could be useful to address this possibility, but extrapolating the regression line in Figure 3A to zero N fixation suggests that N fixation might supply only about 75% of total N, implying a lower EAC. Given these uncertainties, we have not converted our hydrogen-production data to nitrogen-fixation rates. Our measurements seem useful for relative comparisons among rhizobia strains that differ little in EAC, as seems to be the case for our strains, based on the fit of the regression line in Fig. 3A. To calculate absolute N-fixation rates, one would need to measure EAC under conditions that match those for which the actual rates are needed.

Figure 3B challenges the assumption of the “SNA hypothesis” that differences among strains in per-plant benefit in single-strain experiments reflect differences in per-nodule benefit well enough to rank more- and less-beneficial strains. Fourteen of 20 isolates (including overlapping isolates K, L, and M) had a twofold range in whole-plant yield, despite being nearly identical in benefit per integrated nodule mass per plant (Figure 3B). We would therefore not expect differences in performance-based nodule-level sanctions against strain A versus T, for example, even though the difference in per-plant benefits might lead to the mistaken conclusion that strain A is much more beneficial than strain T. In the field, we would expect plants with any mix of those two strains to perform similarly, whereas we would expect seed yield of plants inoculated with a mix of strains A and B (for example) to decrease as the proportion of B increased. Such response-to-proportion tests have been proposed (Oono et al., 2009; Friesen, 2012) as an alternative to single-strain tests. Comparing plant growth with 50% of different focal strains in a background mixture of strains from the same soil would not necessarily require more replicate plants than single-strain experiments.

Results in Fig. 3C are consistent with the hypothesis that hormonal manipulation of plants by rhizobia could have small positive or negative effects on water-use efficiency. Strains E and F had similar seed yields to strains O and C, while using less water. However, we would want to see consistent results from at least one separate experiment before pursuing the mechanisms behind this apparent difference. Field assays of water-use efficiency, like a recent comparison of pea cultivars (Millar et al., 2026), better reflect the effects on nitrogen fixation of vapor-pressure deficit (Monnens et al., 2023), soil water, etc. However, to compare effects of rhizobia strains, rather than legume cultivars, would require controlling nodule occupancy, which might be an insurmountable challenge in the field.

Consistency in strain rankings is arguably essential, either in choosing inocula for agricultural use or for research on phenomena like host-imposed sanctions. We encourage fellow researchers to repeat experiments before drawing broad conclusions. For example, a peptidase gene had contrasting effects between two experimental years (Wendlandt et al., 2022). In one three-year experiment, a storm in one year obscured genetic correlations between nitrogen fixation and seed yield that were apparent in the other two years (Millar et al., 2026). We hope no scientist would publish based on an obviously “bad” year, but more-subtle sources of error may not be so apparent. Our per-nodule strain rankings might be slightly more consistent than rankings that do not correct for differences in nodule mass or number (Figure 4). However, we would still hesitate to claim that strain J would have consistently greater per-nodule benefit than strain P, for example.

Based on Figure 3B, we reject the “SNA hypothesis” that rhizobia strains differ mainly in per-nodule benefits and its implication that per-nodule benefits can be predicted reliably from whole-plant biomass in single-strain experiments. Most basic and applied research comparing strain benefits should either report benefits per nodule and per nodule mass or explain why ignoring differences in nodulation among strains is justified for a particular question. However, our concerns with using vegetative biomass as a proxy for effects on plant fitness or crop yield of soybean were somewhat alleviated by Figure 5, despite conflicting results for other species (Briggs et al., 2026).

Despite the difficulty of displacing indigenous strains via inoculation (Thies et al., 1991), our methods could lead to practical applications. In particular, it may be possible to identify or develop cultivars that selectively enrich soil with more-beneficial strains. Rhizobia can reproduce a millionfold or more in a nodule, so senescing nodules release enough rhizobia to greatly alter soil populations. Five years after soybeans were grown, there were still 25,000 cells/g soil of a marked rhizobia strain, versus 5000 if plants were removed before nodule senescence was complete (Kuykendall, 1989). Both numbers exceed the 10 cells/g needed to out-compete inoculum strains (Thies et al., 1991). Benefits to subsequent crops from cultivars that preferentially support and release more-beneficial strains would be due to positive changes in the composition of rhizobia populations, not increases in total numbers.

In particular, cultivars with stricter sanctions might have beneficial legacy effects on future crops grown in the same soil (Denison & Muller, 2022). Tradeoffs for rhizobia between fixing more nitrogen to avoid host sanctions, versus diverting resources from nitrogen fixation to PHB for future use, could be a constraint. We found that strains that hoarded more PHB while in nodules survived years longer in field soil. However, these differences were only significant after 5 years or more (Muller & Denison, 2024). Even if hoarding less PHB is a necessary (although not sufficient) trait for maximum rhizobia mutualism, PHB-related tradeoffs might not be a serious constraint on sanctions-based legacy benefits in two-year rotations of maize and soybean.

How might cultivars with stricter sanctions be identified? To compare plant responses to nodules differing only in nitrogen fixation, we previously manipulated the partial pressure of nitrogen gas around nodules, all containing the same effective strain. Soybean cultivar “S0066” strongly limited reproduction of rhizobia that fixed nitrogen at 33% or less the rate of control nodules on the same plant (Kiers et al., 2006), but sanctions were not consistently imposed on nodules exposed to 4 kPa nitrogen, which fix at about 50% of control. Manipulating nitrogen gas avoids confounding effects of nitrogen fixation with strain-identity signals. However, low nitrogen concentrations do not perfectly mimic a low-fixing strain. Reducing the nitrogen concentration reduces the nitrogen-fixation rate, but increases the production of hydrogen, which also has a carbon cost to the plant.

Well-defined mutants that fix less nitrogen (Westhoek et al., 2021) might be an ideal tool for identifying cultivars with stricter sanctions against strains that fix less nitrogen, if those differences are consistent over development. But what if benefits to plants depend on differences in the timing of nitrogen fixation (as in Fig. 2A), or on differences in hormonal manipulation (possibly reflected in Fig. 3C)? Indigenous strains might differ in these traits in ways that either complement or undermine differences in nitrogen fixation. Ideally, cultivars would be tested by all three methods: gas treatments, isogenic strains differing in defined traits, and tests using well-characterized field strains. We hope that our methods will contribute to the latter approach.

## Abbreviations

EAC: electron-allocation coefficient
PHB: polyhydroxybutyrate
SNA: specific nodule activity (nitrogen fixation rate per nodule mass)

## Acknowledgments

We thank Mohamed Mohamed and Aaron El-Halawani for plant care and making many of the measurements reported here.

## Author contributions

RFD and NJA designed the experiments and trained RootPainter using root and shoot images, with advice from AGS. KEM isolated the rhizobia and selected the strains used, based on her PHB analyses. RFD designed and built the apparatus and wrote most of the first draft. All authors contributed to the final manuscript.

## Conflict of interest

Gerber files to make the printed-circuit-board instrument shown in Figure 1A will be available as a free, open-source download, but RFD also plans to sell assembled boards at prettygoodpi.org.

## Funding

RFD was supported by funds from the Social Security Administration.

## Data Availability

Data will be available on Data Repository for the University of Minnesota (DRUM) https://hdl.handle.net/11299/166578.

## Notes

### Competing Interest Statement

To facilitate use of our methods, I plan to sell an assembled printed circuit board with the sensors, pumps, etc., in addition to making design files available free for anyone who wants to make their own.

